# Disinfection of Methicillin-Resistant *Staphylococcus aureus*, Vancomycin-resistant *Enterococcus faecium* and *Acinetobacter baumannii* using Klaran WD array system

**DOI:** 10.1101/2020.12.21.423516

**Authors:** Richard M. Mariita, Rajul V. Randive

**Affiliations:** Crystal IS Inc., an Asahi Kasei company, 70 Cohoes Avenue, Green Island, New York, 12183, USA

## Abstract

Healthcare-associated infections (HAIs) are a major burden in healthcare systems. In this study, UVC LEDs emitting radiation from 260-270 nm were evaluated for effectiveness in reducing methicillin-resistant *Staphylococcus aureus* (MRSA), vancomycin-resistant enterococci (VRE), and *Acinetobacter baumannii*. The array has four WD LEDs, each with 70mW placed at 7cm from test organism. With 11.76 mJ/cm^2^, the study obtained 99.99% reduction (log10 reduction factor of 4) against MRSA and VRE. For *A. baumannii*, 9 mJ/cm^2^ obtained 99.999% reduction (log10 reduction factor of 5). These results present scientific evidence on how effective UVC LEDs can be used in the fight against HAIs.

## Introduction

Worldwide, hospital-acquired infections (HAIs) are responsible for extended admissions, increased medical costs, and noticeable morbidity and mortality [1]. Per year, it is approximated that in the US alone, 2 million patients suffer with healthcare-associated infections (HAIs), while nearly 90,000 are estimated to die. The overall direct cost of HAIs to hospitals ranges from US$28 to 45 billion [2]. Microorganisms responsible for HAIs include; Methicillin-Resistant *Staphylococcus aureus* (MRSA), Vancomycin-resistant Enterococcus faecium (VRE) and *Acinetobacter baumannii*. Here, UVC LEDs emitting radiation at 260-270 nm were evaluated for effectiveness in reducing methicillin-resistant *Staphylococcus aureus* (MRSA) ATCC 33592, vancomycin-resistant enterococci (VRE) ATCC 700221, and *Acinetobacter baumannii* ATCC 19606.

*Staphylococcus aureus* is a major pathogen both within hospitals and in the community as healthcare systems in the US and Europe have seen the prevalence of MRSA increase from less than 3% in the early 1980s to rates as high as 40% in the 1990s [3]. In 2017 alone, the US had 119,000 infections and almost 20,000 deaths [4]. MRSA costs about $10 billion a year to treat in the US, averaging about $60,000-$70,000 per patient [5]. MRSA can be transmitted via surgical tools, high touch surfaces, air in surgical units as well via intubation in adult ICU patients, causing pneumonia [3].

VRE is of medical importance, as it is associated with serious multidrug-resistant infections [6]. Moreover, in the US, the emergence of VRE in water is threatening the health of human beings due to antibiotic resistance [6] and the life-threatening nature of some VRE infections [6]. VRE has been isolated from most sites and objects in health care facilities, including medical equipment (ventilator tubing, pumps, wash bowls, automated medication dispensers, intravenous poles), monitoring devices (call bells, electrocardiographic monitors, pulse oximeters, glucose meters, stethoscopes, electronic thermometers, blood pressure cuffs, keyboards, wall-mounted control panels), furniture (telephones, air cushions, headboards, tables, chairs, bed rails), toilet seats, doors, floors and linens [7]. In the US, hospital costs due to VRE infections vary between $9,949 and $77,558 [8].

Globally, *A. baumannii* is one of the most clinically significant multidrug-resistant organisms [9]. *A. baumannii* can be aerosolized, has been found to contaminate trauma ICUs and HVAC systems, especially in locations with high humidity, causing pneumonia and urinary tract and bloodstream infections [9]. In 2017, carbapenem-resistant Acinetobacter caused an estimated 8,500 infections in hospitalized patients and 700 estimated deaths in the United States alone [10] with $281 million in healthcare costs. There are few treatment options, hence the designation of *A. baumannii* as a pathogen of urgent concern and priority [10].

This study explores the disinfection performance of UVC Led array against causative agents responsible for hospital acquired infections (HAIs). Specifically, the study investigated the UVC dose required for inactivation MRSA, VRE and *Acinetobacter baumannii*. The results will be used as baseline in determining the UVC dose required in the eradication of HAIs in healthcare settings to control the causative agents thus minimizing extensive use of antibiotics.

## Materials and methods

### Strain and culture conditions

All strains were stored at −80°C in appropriate culture media using sterile 20% glycerol stock. Working strain cultures were stored at 4°C and propagated before being used in UVC disinfection tests. MRSA and *A. baumannii* strains were propagated in ATCC Medium 3 (Nutrient agar/broth) whereas VRE was propagated in ATCC Medium 44 (Brain heart infusion agar/broth). Cultures were incubated for 18-20 h at 37°C while shaking at 180 rpm. For use, each strain was harvested by centrifugation at 4000 rpm for 10 min. The pellets were washed using 1X PBS three times for 10 min. Between each wash, the supernatant was discarded, and the remaining pellet re-suspended by vortexing. After washing thrice, the pellet was resuspended in 1X PBS used for static dosing study. Agar plates containing irradiated strains were incubated at 37°C for a further 18-20 hrs. Static dosing was performed in three independent replicates and means used in statistics.

### UVC LEDs disinfection

UVC disinfection efficiency of a 4 UVC LED array was evaluated using stationary growth phase bacteria. The array emitted radiation at wavelength between 260 to 270 nm as confirmed using Ocean Optics USB4000 photospectrometer. The distance between the UVC LEDs and agar plate inoculated with bacteria suspension was 7 cm. Disinfection efficacy was assessed after 3, 6, 9, 12 and 15 seconds of irradiation (Table I). Controls were not UVC dosed. Log reduction value (LRV) was calculated using the equation:

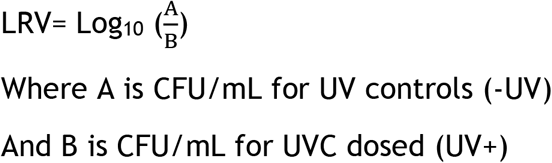

**Table I:** Effectiveness of UVC disinfection against Methicillin-Resistant *Staphylococcus aureus* (MRSA), Vancomycin-resistant Enterococcus faecium (VRE) and *Acinetobacter baumannii* at 7cm using an array with 4 UVC LEDs. Experiment performed in triplicate and data expressed as mean.

| Strain | UVC dose (mJ/cm<sup>2</sup>) | Irradiation time (Sec) | Control CFU/mL (-UVC) | Test CFU/mL (+UVC) | LRV |
| --- | --- | --- | --- | --- | --- |
| VRE | 3.15 | 3 sec | 1.00E+09 | 1.11E+07 | 1.96 |
| VRE | 6.18 | 6sec | 1.00E+09 | 5.43E+06 | 2.26 |
| VRE | 9.00 | 9 sec | 1.00E+09 | 6.67E+05 | 3.18 |
| VRE | 11.76 | 12 sec | 1.00E+09 | 5.57E+04 | 4.25 |
| VRE | 16.5 | 15 sec | 1.00E+09 | No growth | - |
| MRSA | 3.15 | 3 sec | 1.61E+08 | 1.10E+07 | 1.17 |
| MRSA | 6.18 | 6sec | 1.61E+08 | 1.00E+06 | 2.21 |
| MRSA | 9.00 | 9 sec | 1.61E+08 | 1.33E+05 | 3.08 |
| MRSA | 11.76 | 12 sec | 1.61E+08 | 1.00E+04 | 4.21 |
| MRSA | 16.5 | 15 sec | 1.61E+08 | No growth | - |
| *A. baumannii* | 3.15 | 3 sec | 2.49E+07 | 3.55E+03 | 3.85 |
| *A. baumannii* | 6.18 | 6sec | 2.49E+07 | 2.30E+02 | 5.03 |
| *A. baumannii* | 9.00 | 9 sec | 2.49E+07 | 8.67E+01 | 5.46 |
| *A. baumannii* | 11.76 | 12 sec | 2.49E+07 | 2.53E+01 | 5.99 |
| *A. baumannii* | 16.5 | 15 sec | 2.49E+07 | 3.33E+00 | 6.87 |

To determine strength or relationship between irradiation time and disinfection efficacy against each test strain, linear regression analysis was performed using GraphPad Software (https://www.graphpad.com/).

## Results and discussion

Table I shows that the UVC LED array is very effective against study test strains. In all cases, 11.76 mJ/cm^2^ was enough to obtain > LRV4 (99.99% reduction). Additionally, the study revealed that *A. baumannii* was more susceptible to UVC radiation than MRSA and VRE strains (Figure 1).

**Figure 1:**
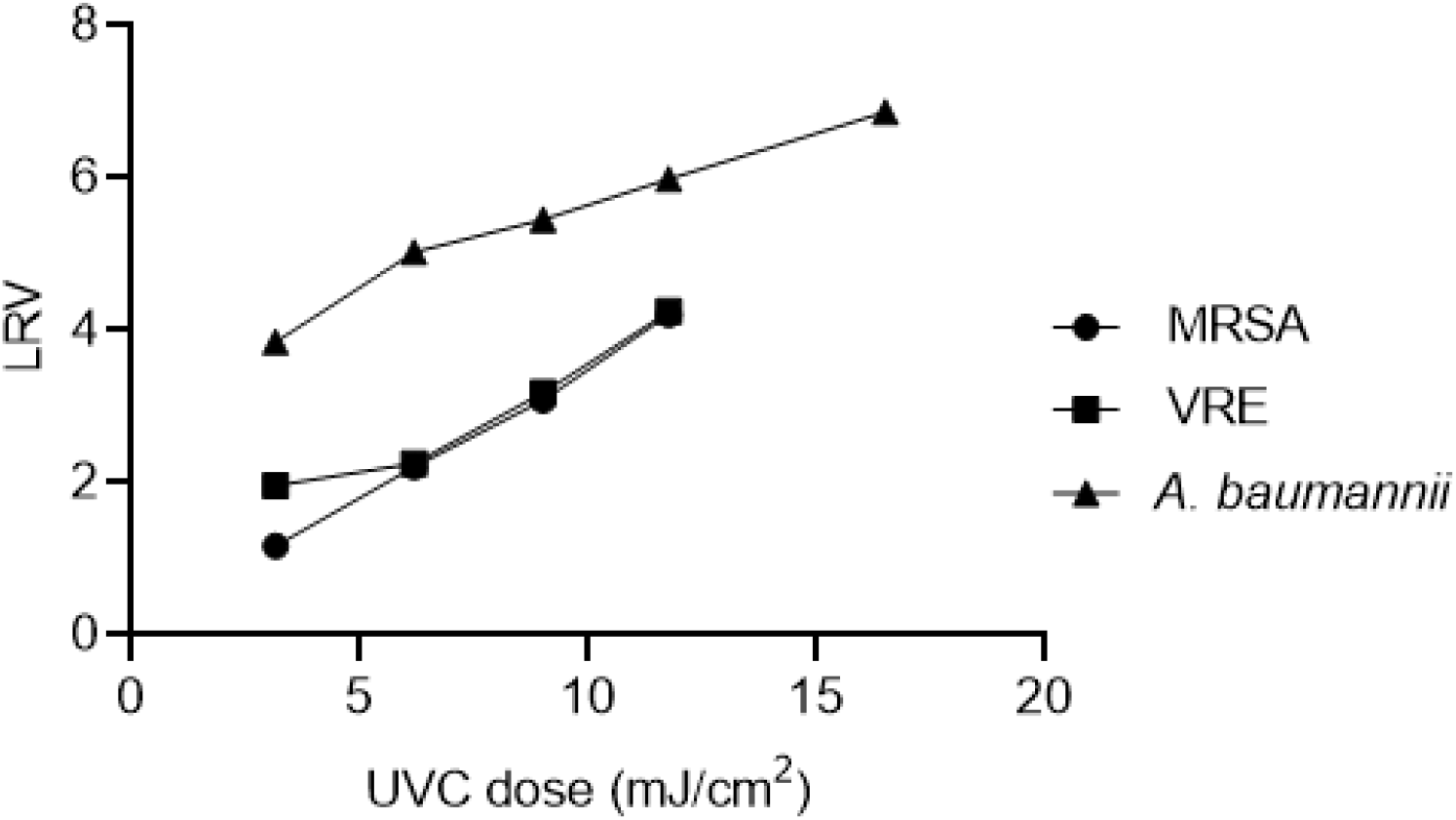
Disinfection efficacy of UVC against key pathogens associated with HAIs. *A. baumannii* was more sensitive against UVC compared to MRSA and VRE

Linear regression analysis at 95% for MRSA (R^2^=0.9969, *p*=0.0016), *A. baumannii* (R^2^=0.9681, *p*=0.0024) and VRE (0.9403, *p*=0.0303) displayed strong association between irradiation time and disinfection efficacy. Based on the findings, it is proposed that a further feasibility study in areas such as in Intensive care and surgical units as well as other high touch surfaces be undertaken.

### Conclusion

The results from this study suggest that arrays of Klaran WD UVC LEDs emitting radiation between 260-270 nm can provide effective and rapid decontamination of HAIs. We intend to carry out *in situ* tests to assess usability and relative performance.

## Acknowledgments

Authors thank Michelle Lottridge, Amy Miller and Chris Scully for their technical help and review of manuscript.

